# Physiological re-replication during human stem cell differentiation

**DOI:** 10.64898/2026.02.27.708451

**Authors:** Marie Minet, Amila Beganovic, Shusruto Rishik, Elisa Michaeli, Daniela Yildiz, Georges P Schmartz, Paula E. Schwarz, Melina Schäfer, Tanja Tänzer, Magali Cucchiarini, Nicole Ludwig, Andreas Keller, Eckart Meese, Ulrike Fischer

**Affiliations:** Institute of Human Genetics, Saarland University, Homburg, 66421, Germany; Chair of Clinical Bioinformatics, Saarland University, Saarbrücken, 66123, Germany; Chair of Molecular Pharmacology, Center for Molecular Signaling (PZMS), Saarland University, Homburg, 66421, Germany; Center of Experimental Orthopedics, Saarland University Medical Center, Homburg, 66421, Germany; NGS Sequencing Facility, Medical Faculty, Saarland University, Homburg, 66421, Germany

## Abstract

During defined developmental windows in *Drosophila*, controlled re-replication generates physiological gene amplification. Although gene amplification has also been observed during human stem cell differentiation, re-replication in human cells has largely been linked to tumor-associated genome instability. Here, we demonstrate that re-replication likewise operates as a physiological mechanism in human stem cells. Using Rerep-Seq and DNA fiber-combing, we identify distinct phases of re-replication during the differentiation of human myoblasts into myotubes and during the lineage commitment of mesenchymal stem cells toward adipogenic, osteogenic, chondrogenic, and neuronal fates. In all differentiation systems examined, re-replication occurred within defined temporal windows. FACS-isolated re-replicating cells exhibited elevated gene expression using RNA-Seq specifically within re-replicated genomic regions. Moreover, re-replicated DNA was detected as extranuclear DNA. These findings support a model in which cells that do not undergo re-replication, and thus avoid increased chromosomal instability, may nonetheless boost the expression of differentiation-relevant genes by acquiring re-replicated DNA released from neighboring re-replicating cells. We propose that human stem cells exploit an evolutionarily conserved re-replication mechanism to transiently increase gene copy number and thereby meet the heightened protein demands associated with differentiation.

## INTRODUCTION

Chorion gene amplification in *Drosophila* eggshell cells has been extensively studied for nearly four decades. Osheim (1988) and Alexander (2015) suggested an underlying mechanism whereby cell cycle control, which typically limits replication to once per cell cycle, is bypassed to enable endoreplication (1, 2). While multiple origin-firing leading to re-replication has been well-studied in *Drosophila*, less data is available on re-replication in mammalian cells. Here, re-replication events have primarily been described in pathological contexts such as tumors, genome instability and as inducers of gene amplification (3). Pathways that inhibit re-replication have also been linked to prevention of tumorigenesis (4). The amplification has only more recently been shown in a physiological context during the differentiation of stem cells (5, 6) raising the question of whether physiological amplification processes in human cells are also based on re-replication events.

Genes that have been described in various species in connection with the regulation of replication offer starting points for addressing this question. The proteins CDT1 and CDC6 are vital for origin licensing and DNA replication initiation (7). CDT1 is primarily repressed after replication begins to prevent further origin licensing and re-replication (8, 9). Mammalian cells inactivate CDT1 through mechanisms such as ubiquitination, degradation, and binding by Geminin (GMNN) (4). Both *CDT1* overexpression and *GMNN* depletion can lead to re-replication (10, 11). In addition, a decreased binding of CDT1 caused by *GMNN* mutations probably induces re-replication and cell cycle disturbance (12).

In addition to these biological targets, there are two methodological approaches to address the question of re-replication in human stem cells. Fiber-combing allows for the visualization of multiple origin firings but does not specify which DNA regions are re-replicated. This can be achieved by a technique called Rerep-Seq, which connects re-replication detection with sequence information (13).

By analyzing the differentiation of human myoblasts and human mesenchymal stem cells (hMSCs), we address three key questions: I) Can re-replication occur as a potential mechanism for gene amplification in human stem cells, like the process observed in *Drosophila* eggshell development? II) Which specific regions of DNA are re-replicated? III) Are genes located in re-replicated DNA regions subject to overexpression?

Furthermore, it is unclear how normal human cells manage to survive with extra DNA while mitigating the risks associated with genome instability, a challenge that *Drosophila* eggshell cells do not face. To explore this issue, we investigate two further questions: I) Can we detect extranuclear re-replicated DNA? II) Is re-replication detectable on both strands of DNA?

## MATERIAL AND METHODS

### Cell Culture and Differentiation

HSkM (Human Skeletal Myoblast) cells derived from primary normal human skeletal myoblasts were obtained from Thermo Fischer Scientific (Waltham, MA, USA). These cells undergo differentiation to myotubes within 48h. HSkM cells were cultivated in Dulbecco’s Modified Eagle Medium (DMEM) supplemented with 2% Horse Serum to induce differentiation.

Human BMSCs (Bone marrow Mesenchymal Stem Cells) were obtained from PromoCell GmbH (Heidelberg, Germany). These cells were approved and certified by the supplier with immune staining and flow cytometric analyses. The cells were seeded at a density of 100,000/25 cm^2^ flask with Mesenchymal Stem cell Growth Medium (PromoCell) and expanded for 1 passage before induction of differentiation. Induction of differentiation was for adipogenic, osteogenic and neuron differentiation on fibronectin coated cell culture flasks and for chondrocyte differentiation on uncoated cell culture ware. Recommended densities were 70-80% for adipocytes, 100% for osteoblasts, 60% for neurons and 80-90% for chondrocytes, each cultivated with corresponding differentiation media from the supplier (PromoCell). Although we investigated only differentiation towards adipocytes, osteoblasts, chondrocytes and neurons until 72h in this study, we confirmed the differentiation in biological replicate experiments after 10d for adipocytes (lipid vacuole), 12d for osteoblasts (anti-osteocalcin), 7d for chondrocytes (toluidine blue stain) and 2d for neurons (mRNA expression of synapse maturation and neuron markers (14)). For laser microdissection, hMSC (human Mesenchymal Stem Cells) cells were grown on glass slides with fibronectin (adipocyte and osteoblast differentiation) and without fibronectin (chondrocyte differentiation) as described above.

### Rerep-Seq

For detection of re-replication the original protocol by Menzel and colleagues describes the addition of the thymidine-analogue BrdU to the culture medium (13). Normal replication results in double-stranded DNA with one BrdU-incorporated strand and one parental strand, while re-replication incorporates BrdU into both strands. After UVA treatment with Hoechst and processing with AP1 and UDG enzymes, normal replication results in strand breaks only in the new strand at BrdU sites, whereas re-replication causes breaks in both strands, leading to DNA fragmentation. We deviated from the Menzel protocol by using 10µg of DNA for UVA and enzyme treatment and we added a control experiment without BrdU for each differentiation. DNA from harvested cells was isolated by NaCl/chloroform extraction after RNase treatment. In total 10µg DNA from cells with BrdU and without BrdU (control experiment) treatment were UVA treated as described (13) and separated by gel electrophoresis. Fragments were excised from >100bp to 1.5kb, reisolated and used for next generation sequencing analysis using MGIEasy Universal DNA Library Prep Kit and HotMPS High-throughput Sequencing Set G400 HM FCL PE100 (MGI Tech, Shenzhen, China). All data was processed using the nf-core/chipseq pipeline, which is build using Nextflow (15). We selected the nf-core/chipseq pipeline not only for its comprehensive quality control processes, adapter trimming, and alignment but also because it aligns with our primary interest in DNA coverage profiles, including normalized bigWig files and MACS2 for peak calling. This pipeline is particularly well-suited to our needs due to its robust implementation to generate precisely these coverage profiles. To identify significant regions of enrichment, we adjusted the p-value threshold to 0.1 to ensure appropriate sensitivity for our analysis. By setting this threshold, we ensured the capture of potentially biological relevant peaks.

### Thymidine-analogue treatment

For fiber-combing analysis, cells were differentiated as described above and treated with thymidine-analogues as follows. IdU (Iodo-deoxyuridine) was added to differentiating cells with a final concentration of 60μM for 1.5h, after which the cells were briefly washed with PBS, given fresh differentiation media for 15 minutes, and given the second pulse of CldU (Chloro-deoxyuridine) for 45 minutes with a final concentration of 500 µM. After all pulse labeling steps, the cells were harvested using Accutase, resuspended in PBS and processed further. For Rerep-seq analysis, cells were differentiated as described above and BrdU was added to differentiation media with a final concentration of 48µM for 8h (chondrocyte differentiation and neuron differentiation), 12h (osteoblast and myoblast differentiation) and 14h (adipocyte differentiation). For FACS and laser microdissection, cells were differentiated as described above and EdU was added to differentiation media with a final concentration of 90µM and 120µM respectively.

### Fiber Preparation for Molecular Combing

Harvested cells from culture (adipocyte, osteoblast and neuron differentiation) were diluted and further processed as described in the manual of FiberPrep Kit (Genomic Vision, Bagneux, France). The resulting DNA suspension was transferred to a combing reservoir inserted into the FiberComb Molecular Combing Device, along with 1-2 silanized coverslips (Genomic Vision, Bagneux, France) per reservoir. The coverslips were then mechanically inserted into the DNA suspension and pulled out at a constant rate. This resulted in DNA fibers adhering to the silanized surface with a constant measurement of 2kb/μm. Once the DNA was combed onto the coverslips, they were denatured at 85 °C for 5 minutes.

### Fluorescent Immunostaining of Combed Fibers

Coverslips were blocked with 10 % goat serum diluted in PBS for 30 minutes at 37°C. This is followed by a mouse-anti-BrdU-antibody (for IdU detection) and a rat-anti-BrdU-antibody (for CldU detection) diluted 1:10 in 10 % goat serum, and incubated for 1 hour in a humid chamber at 37°C. The coverslips were subsequently washed in PBS and incubated in stringency buffer for 10 minutes to enhance the specificity of primary antibody binding. Then, the corresponding secondary antibody solutions goat-anti-mouse IgG (Alexa Fluor Plus 594) and goat-anti-rat IgG (Alexa Fluor Plus 405), diluted in 10 % goat serum, were placed on the coverslips for 30 minutes in a humid chamber at room temperature. The coverslips were then washed again, and counterstained with YOYO. All coverslips were individually analyzed using Zeiss microscope with ZEN (blue edition) software. Only for chondrocyte differentiation combing and immunostaining were performed according to the EasyComb procedures (Genomic Vision, Bagneux, France). Coverslips were scanned with the FiberVision® scanner and images were analyzed using Genomic Vision FiberStudio® software (Genomic Vision, Bagneux, France). Intact IdU (green) and CldU (red) replication tracks, flanked by counterstaining (blue), were selected and used for further validation.

### Cell sorting

hMSCs were differentiated towards adipocytes, osteoblasts, and neurons as described above. In a first experiment, the thymidine-analogue EdU was added to differentiation media for 0-15h (neuron differentiation) and 6-20h (adipocyte differentiation) and cells were harvested immediately for fixation. In a second experiment, EdU thymidine-analogue was added to differentiation media for 0-15h (neuron differentiation) and 6-20h (adipocyte and osteoblast differentiation) and afterwards fresh differentiation media was added, and the cells were harvested after 24h and 48h for fixation. Next, cells were stained with Click-iT^TM^ EdU Alexa Fluor^TM^ 594 Imaging Kit including DNA counterstaining with Hoechst33342. Fluorescence signal detection and sorting were performed using the Sony SH800 (Sony, Berlin, Germany) and FACS Aria Fusion (BD, Heidelberg, Germany). The gating strategy is indicated in Suppl. Figure S2. Data were visualized with FlowJo 10.6.2 software (Tree Star, Inc., Ashland, OR, USA).

### RNA-seq

RNA was isolated using miRNAeasy Kit (Qiagen, Hilden, Germany) and used for next generation sequencing. Whole transcriptome sequencing was performed using MGIEasy Library Prep Set and DNBSEQ-G400RS High-throughput Sequencing Set (FCL PE100) (MGI Tech, Shenzhen, China) with 100ng total RNA input. In short, rRNA was depleted using the MGIEasy rRNA depletion kit and remaining RNAs were fragmented to 150bp size using heat fragmentation. After reverse transcription, end repair and A tailing, barcoded adaptors were ligated to the cDNA at the 3‘and 5’ends. Ligation products were PCR amplified for 12 cycles and purified using magnetic beads. All 16 samples were pooled into one library and circularized using a specific oligo sequence complementary to sequences in both the 3‘and 5’adaptors, remaining linear DNA was digested. Subsequently, DNA nanoballs (DNBs) were generated using rolling circle amplification. DNBs were loaded onto the flowcell and sequenced with the DNBSEQ-G400RS High-throughput Sequencing Set PE100 generating paired end 100bp reads on the DNBSEQ-G400RS sequencer (MGI Tech, Shenzhen, China). Paired-end fastq files were processed using the mRNA module of SnakePipes 3.0.0 (16). The reads were aligned using STAR using default settings provided by snakepipes (17) against GRCh38 (18) in order to generate bam and bai files. Quality control checks were carried out using multiQC (19).

### Laser microdissection of extranuclear DNA

Differentiated hMSC on glass slides were stained by Edu-Click-iT^TM^-reaction with Alexa Fluor-594 and counterstained with Hoechst33342. Extranuclear DNA clusters or spots were identified using RoboSoftware 4.9 and LCM/AxioVision SE64 Rel. 4.9.1 Software with 20x magnification. Up to 200 areas were selected and sequentially laser-microdissected with PALM MicroBeam Rel. 4.2 system into AdhesiveCap 500 clear tubes (Zeiss, Germany). DNA from each tube was isolated using QIAmp DNA Micro Kit according to the instructions for laser-microdissected material. For each differentiation experiment 6-8 DNA isolations were pooled, vacuum dried and resuspended in TE for sequencing (NGS).

## RESULTS

To analyze re-replication, we use two methods including Rerep-Seq and fiber-combing that complement each other. Each method addresses specific aspects of re-replication. While fiber-combing provides visual evidence of replicated DNA strands at the single-molecule level, the Rerep-Seq method allows for the genomic localization of such events. Since Rerep-Seq is a bulk analysis, it does not examine individual molecules and since Rerep-Seq involves DNA fragmentation, it prevents meaningful lengths determination of re-replicated regions. A summary of our experimental strategy (A) and a timeline overview on Rerep-seq experiments, thymidine-analogue pulse for fiber-combing (B) is given in Figure 1.

**Figure 1:**
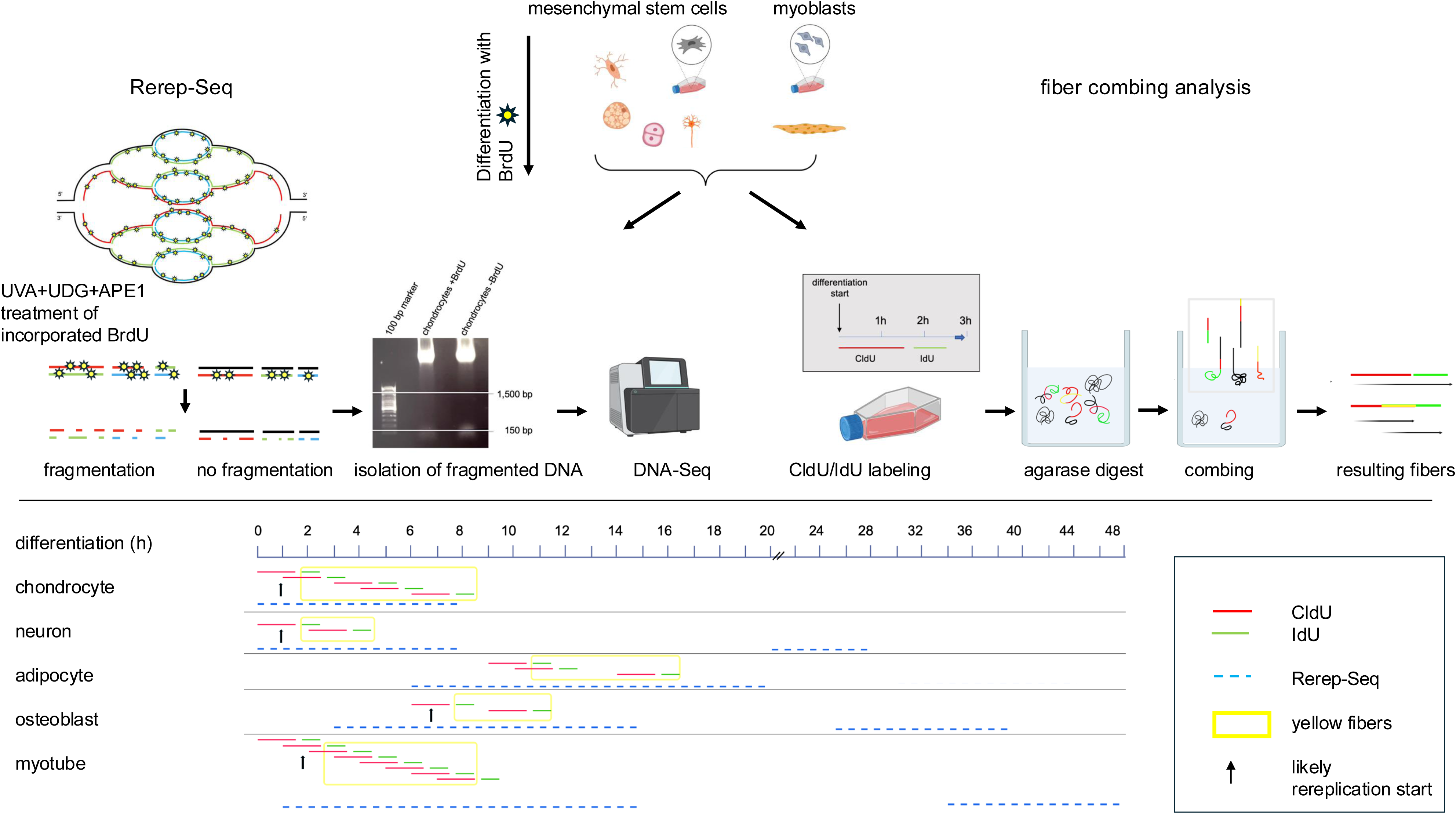
Overview and time scale of re-replications experiments Mesenchymal stem cells were differentiated into osteoblasts, adipocytes, chondrocytes, and neurons, while myoblasts were differentiated into myotubes. Two experimental strategies were applied to detect re-replication during differentiation (A). For Rerep-seq the thymidine-analogue BrdU was added to the culture media. In the schematic representation of the re-replication bubble, yellow asterisks indicate BrdU integration. The black DNA strand represents parental DNA, the red strand corresponds to DNA from the first round of replication, the green strand to DNA from the second round, and the turquoise strand to DNA from the third round of replication. As described by Menzel et al. (13), UVA photolyzes incorporated BrdU converting it to Uracil. Subsequently, UDG excises the Uracil, generating an abasic site, while AP1 introduces single strand breaks at these positions. A representative agarose gel image demonstrates the effects of UVA, UDG, and AP1 treatment on DNA from chondrogenic differentiated hMSCs cultured with and without BrdU supplementation. For fiber-combing thymidine-analogues IdU and CldU were subsequently added to culture media. High molecular weight DNA was used for fiber-combing and further evaluated for fibers with simultaneous IdU and CldU incorporation. Time scale shows an overview on start and end points of all differentiation experiments (B). Yellow rectangles highlight timeframes where re-replication was detected with fiber-combing. Black arrows point on likely re-replication start points during the differentiation process. The figure was created using BioRender.

### Genome-wide re-replication during differentiation detected by Rerep-Seq

For Rerep-Seq analysis we selected timeframes with a high probability of occurrence of re-replication estimated from *CDC6* and *CDT1* mRNA expression results. This strategy was successfully applied recently during human myoblast differentiation where a high *CDC6* expression during first 12h and increased *CDT1* expression during first 3h of myoblast differentiation indicated a timeframe with high probability of re-replication that could be confirmed with fiber-combing experiments (20). Timeframes with low probability of occurrence of re-replication were also included to exclude methodological artifacts. Timeframes with a high probability of re-replication were determined using RNA-seq for osteoblast, adipocyte, neuron and chondrocyte differentiation during the first 24h as shown in supplFigure 1. For osteoblast-and adipocyte-differentiation *CDT1* mRNA expression revealed first a decrease of *CDT1* expression and around 3h for osteoblast and 6h for adipocyte stabilized *CDT1* and *CDC6* expression and a further decrease after 12h until 24h post differentiation induction. For neuron and chondrocyte differentiation expression pattern is different with a high expression of *CDT1* and *CDC6* at the beginning of neuron differentiation and an increase of *CDC6* mRNA expression 3h after neuron and chondrocyte differentiation induction. To find timeframes with low probability of re-replication we used Click-it^TM^-EdU-Imaging-assay to survey incorporation of the thymidine-analogue EdU. During the first 24h EdU is incorporated in 41% of neuron, 60% of adipocyte, 74% of osteoblast and 66% of chondrocyte differentiating cells. Only during neuron differentiation addition of EdU after 2d revealed no further incorporation. During adipocyte, osteoblast and chondrocyte differentiation EdU incorporation after 2d was detectable in 3%, 8% and 10% of the cells and was reduced to 0.1%, 1% and 0.2% of the cells after 6d of differentiation.

In detail, timeframes with high probability of re-replication included, 3-15h osteoblast differentiation, 6-20h adipocyte differentiation, 0-8h chondrocyte and neuron differentiation and 1-15h for HSkM/myotube differentiation. Since EdU incorporation was still detectable until 2d we included timeframes of 25-39h for osteoblast and 30-44h for adipocyte differentiation. Timeframes with low probability of re-replication included, 0-3h for osteoblast and 0-6h for adipocyte differentiation and 72-84h for osteoblast, neuron and adipocyte differentiation. Since HSkM cells were supposed to differentiate to myotubes within 48h as stated by the supplier we included a timeframe of 34-48h for low probability of re-replication.

To distinguish between re-replication-induced fragmentation and random fragmentation, we performed control experiments without BrdU using the same timelines. The fragments were separated by gel electrophoresis as exemplarily shown for osteoblast differentiation in Figure 2A. Fragments of expected sizes (>100bp-1.5kb) were visible best in lane 3-15h BrdU osteoblast. Fragments in lanes 25-39h BrdU osteoblast differentiation were less and mostly around 100bp but sufficient for isolation and sequencing. Experiments 0-3h osteoblast and 72-84h osteoblast differentiation revealed no fragments (>100bp-1.5kb) sufficient for isolation and sequencing.

**Figure 2:**
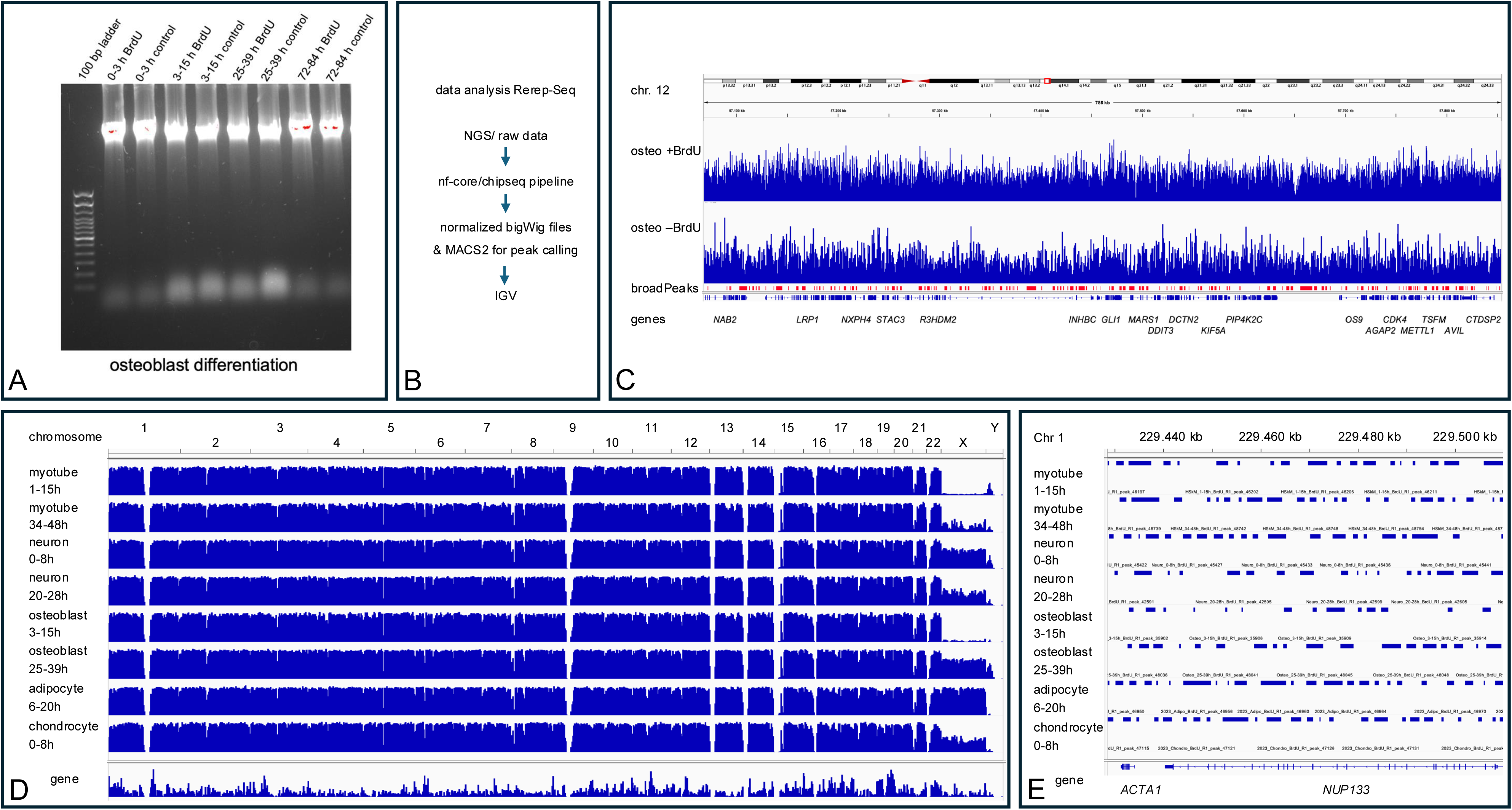
Rerep-Seq results Fragmented DNA from Rerep-seq experiments during four time-windows during osteoblast differentiation were shown in A. Following isolation of fragments of >100bp to 1.2kb NGS was done and a data analysis strategy for Rerep-seq is summarized in B. Exemplarily for chromosome 12 region results of Rerep-Seq data (normalized bigwig files) for osteogenic differentiation with and without BrdU were shown using Integrative Genome Viewer (IGV) (C). A summary of the genome wide results of MACS2 peak calling analysis as broadPeaks on all investigated timeframes of differentiations were displayed in D. In addition, a detailed overview on results of Rerep-Seq analysis using IGV is given for chromosome 1 region including genes *ACTA1 and NUP133*.

As summarized in Figure 2B, raw sequencing data files from experiments with and without BrdU are used as an input for the nf-core/chipseq-pipeline that allows data visualization in the Integrative-Genomics-Viewer (IGV) (21). MACS2 as part of the pipeline creates broadPeaks for all chromosomes with peak height as the signal strength of the data at a particular location. BroadPeaks thereby detected genomic regions with significant enrichment of sequencing reads indicating re-replicated chromosome regions in experiments with BrdU versus without BrdU as exemplarily shown for chromosome 12 region in Figure 2C.

In general, we identified broadPeaks of re-replicated DNA throughout the genome as shown in Figure 2D for all investigated differentiations/timeframes. An exemplarily chosen detailed view of chromosome 1 including genes *ACTA1* and *NUP133* revealed differences in frequency and length of detected broadPeaks between early neuron differentiation and late neuron differentiation within the latter less and shorter broadPeak regions (Figure 2E). Whereas during osteoblast differentiation less and shorter broadPeak regions were detected in the early timeframe of differentiation. Overall, the broadPeak regions in the early and late timeframes showed only partial overlap, yet together they covered the entire region to achieve complete gene coverage (Figure 2E).

### Re-replication during differentiation detected by Fiber-combing

To analyze re-replication using fiber-combing, differentiating cells were pulse-treated with the thymidine analog IdU for 1.5h, followed by a 15-minute incubation in media without thymidine analogs, and then treated with CldU for 45min. We started the pulse treatment so that a two-color staining that indicates re-replication falls within the above detected timeframes for re-replication. That means that all pulse treatments were started and finished, before the end of the timeframes. In detail, we started pulse treatment during neuron differentiation at 0h and 2h after differentiation induction, during osteoblast differentiation at 6h and 9h, during adipocyte differentiation at 9h, 10h and 14h, and during chondrocyte differentiation at 0h, 4h, and 6h. At all analyzed time points fiber-combing detected re-replication events as shown in Figure 3 and Figure 4, the latter one summarizing the fiber lengths.

**Figure 3:**
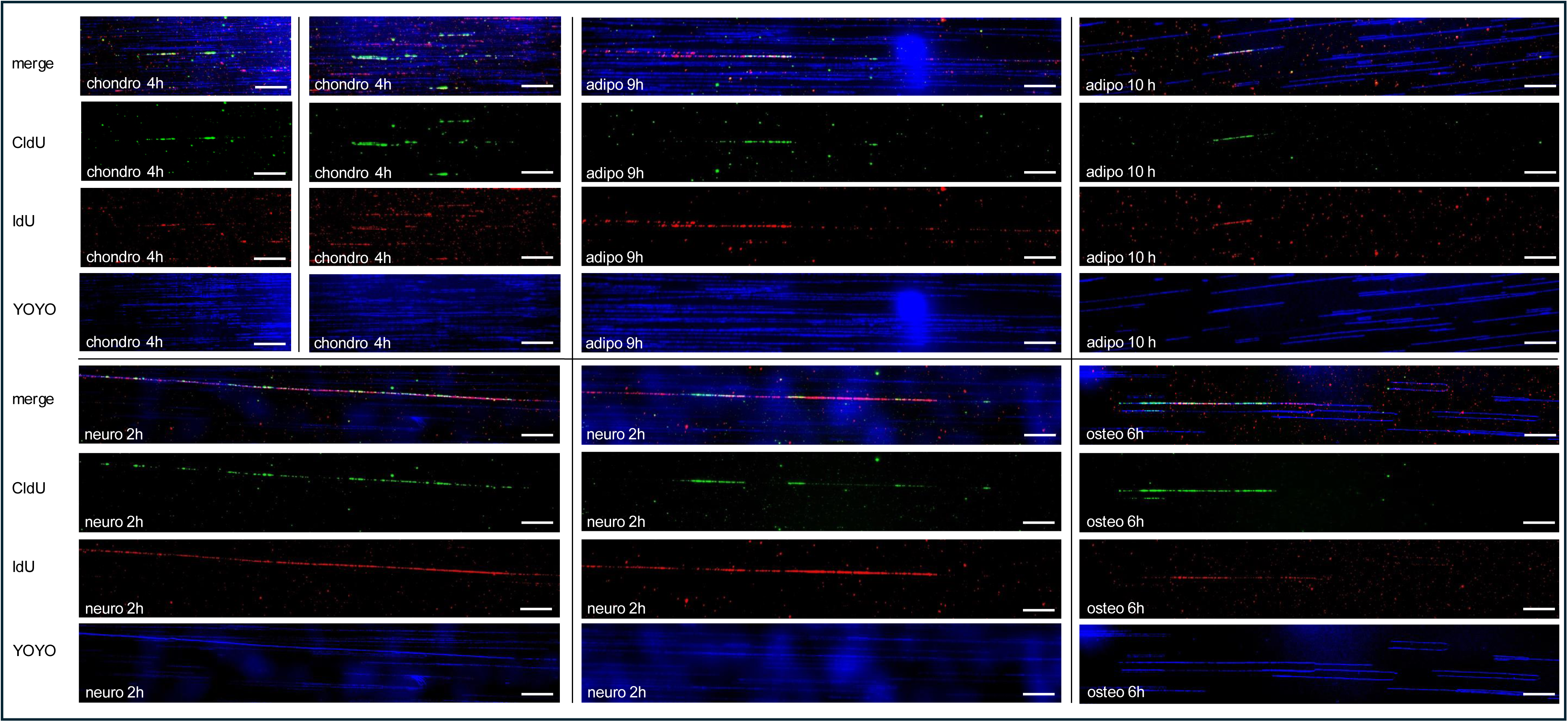
Re-replication analysis using fiber-combing Representative examples of fiber-combing experiments were shown for 4h chondrogenic differentiation (4h chondro), for 9h and 10h adipogenic differentiation (9h adipo and 10h adipo), for 2h neuronal differentiation (2h neuro) and for 6h osteogenic differentiation (6h osteo). For each fiber individual fluorescence and merged staining pictures were presented. DNA is shown in blue (false color YOYO-stain) and thymidine-analogue detection is shown in red for IdU and green for CldU. Re-replicated fiber tracks appear as yellow fluorescence on merged staining pictures. Scale bars represent 20 µm = 40kb.

**Figure 4:**
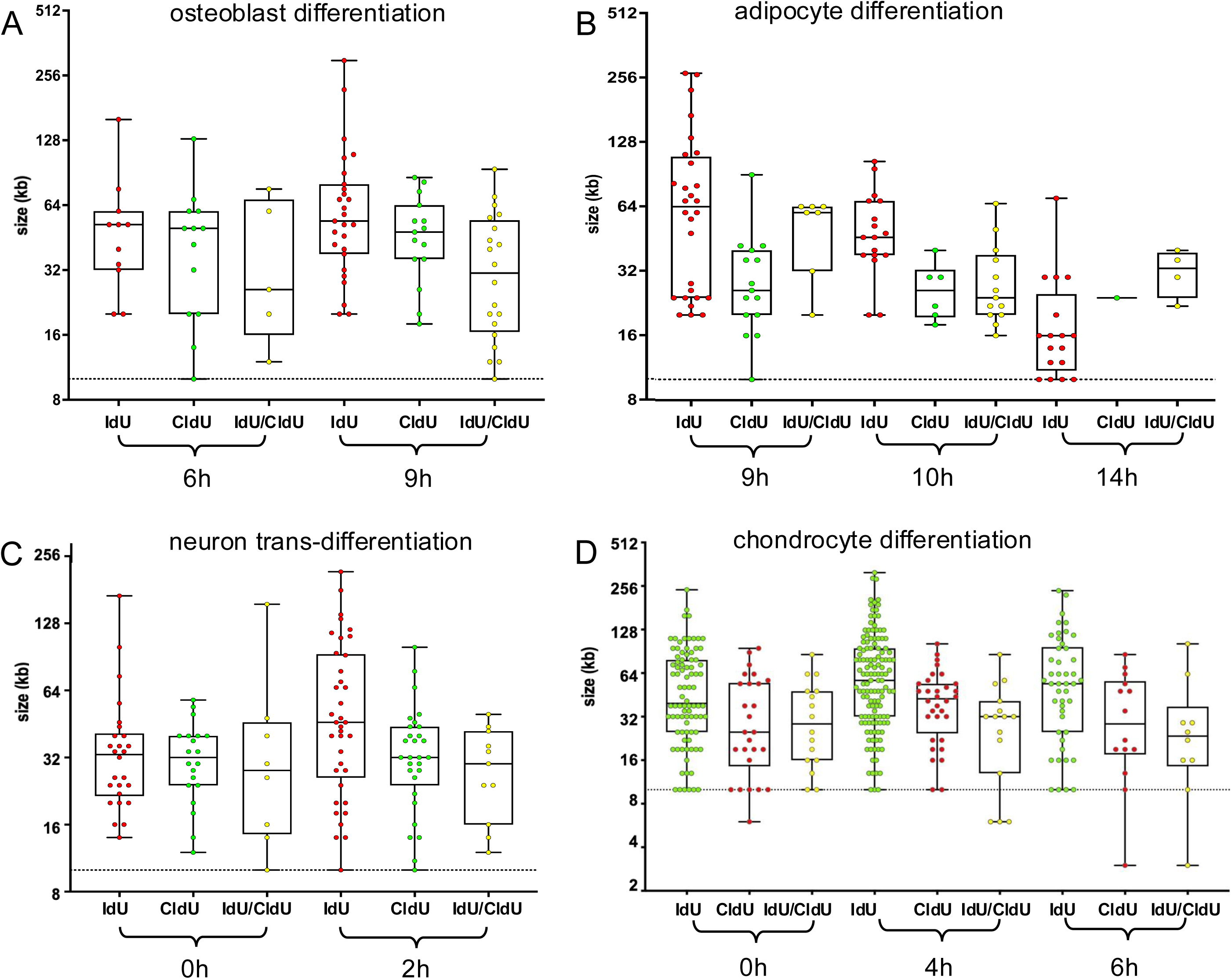
Distribution and length of thymidine-analogue integration Fibers of hMSC cells, which differentiated for 6h and 9h towards osteoblasts (A), 9h, 10h and 14 h towards adipocytes (B), 0h and 2h towards neurons (C), 0h, 4h and 6h towards chondrocytes (D) were analyzed for integrated thymidine-analogues. Results were displayed as dots and represent the length of uninterrupted thymidine-analogue integration of the respective analyzed fiber. The boxes indicate the 2nd and 3rd quartiles, and the whiskers represent the minimum and maximum values. The IdU fiber lengths are shown in red, the CldU fiber lengths are shown in green, and the simultaneous IdU/CldU fiber lengths are shown in yellow (A-C) and IdU fiber lengths are shown in green, the CldU fiber lengths are shown in red, and the simultaneous IdU/CldU fiber lengths are shown in yellow (D). Dotted lines represent the 10kb threshold. The Y axis shows the length of the colored fibers, and the X axis shows the thymidine-analogues used in the different experiments.

Re-replication is detected by yellow tracks in the merged view resulting from simultaneous IdU and CldU incorporation in the combed DNA fibers. Figure 3 shows examples of re-replication detected in chondrocytes at 4h differentiation, in adipocytes at 9h and 10h of differentiation, in osteoblasts at 6h of differentiation, and in neurons at 2h of differentiation. Re-replication analysis during neuron differentiation revealed multiple replication restarts. As previously reported by Dorn et al., short yellow tracks (<2.5kb) were observed at the junction of the first and second pulses, indicating activity from a single replication fork during both pulses. They established a threshold of 1µm (2.5kb) for identifying true yellow tracks, characterized by significant overlap of red and green signals, which indicates re-replication (10). In our study, we focused exclusively on yellow fiber-tracks longer than 10kb as indicators of re-replication events to reduce false positives arising from the persistence of the first thymidine-analogue label.

The lengths and numbers of re-replicated regions varied across differentiation experiments (Figure 4). Notably, more and longer DNA fiber-tracks with incorporated IdU (red) were observed after 9h compared to 6h of osteoblast differentiation (Figure 4A). Similarly, the number of yellow fiber-tracks from simultaneous IdU and CldU incorporation increased after 9h versus 6h. During adipocyte differentiation, both the size and number of yellow fiber-tracks decreased over time (Figure 4B). In neuron differentiation, these metrics remained largely stable between 0h and 2h (Figure 4C), and in chondrocyte differentiation number of yellow fiber-tracks decreased between 0h and 6h (Figure 4D). For each differentiation experiment several hundred fibers were analyzed and revealed an overall frequency of yellow fiber-tracks between 0.5% (14h adipocyte differentiation) and 2.1% (6h chondrocyte differentiation). In addition, fiber length analysis revealed that at 6h osteoblast and 0h neuron differentiation red (IdU) and green (CldU) fibers showed a similar median length, although IdU supplementation was twice as long as CldU supplementation. We conclude that first replication started 0.75h later than the start of first thymidine-analogue pulse.

In summary, both bulk analysis by Re-Rep-Seq and single molecule analysis by fiber-combing demonstrate re-replication during hMSC and myoblast differentiation.

### Expression analysis of replicating versus not-replicating cells

Bulk expression analysis of differentiating hMSCs towards neurons showed no overexpression of amplified genes (14). This could be because only a small portion of differentiating cells carry amplified genes (5, 22). Therefore, we enriched actively replicating cells prior to mRNA expression analysis. To this end, differentiating cells were grown in the presence of EdU, which is incorporated during replication.

In detail, hMSCs were differentiated towards adipocytes with Edu added during the timeframe 6-20h, towards neurons with EdU added from 0-15h and towards osteoblasts with EdU added from 6-20h. FACS sorting was done at 0h after the EdU supplementation of differentiating cells towards adipocytes and neurons, and at 24h and 48h after EdU supplementation of differentiating cells towards adipocytes, neurons and osteoblasts. We used Alexa-594 fluorescence-dye (red) for EdU and Hoechst33342 (blue) for total DNA staining. FACS of cells analyzed directly after the EdU supplementation window revealed 30% EdU-positive (red) cells and 70% Edu-negative (blue) cells during neuron differentiation and 36% EdU-positive cells and 64% EdU-negative cells during adipocyte differentiation. FACS sorting criteria and results are summarized in supplementary Figure S2.

We conducted a differential expression analysis comparing replicating (EdU-positive) and non-replicating (EdU-negative) cells, with an emphasis on re-replicated DNA-regions indicated by broadPeaks as shown in Figure 5. Leptin gene *LEP* showed an enrichment of broadPeaks, and increased RNA read counts in replicating (EdU-positive) cells at 24h and 48h after EdU addition, compared to non-replicating (EdU-negative) cells during differentiation into adipocytes (Figure 5A). *MDM2* gene showed an enrichment of broadPeaks, and increased RNA read counts in replicating (EdU-positive) cells at 24h and 48h after EdU addition during differentiation into adipocytes and osteoblasts (Figure 5B and D). *SYP* gene showed an enrichment of broadPeaks, and increased RNA read counts in replicating (EdU-positive) cells at 24h after EdU addition during differentiation into neurons (Figure 5C). Interestingly several genomic regions all over the genome revealed no broadPeaks and re-replication and genes localized to these regions revealed similar RNA read counts in replicating (EdU-positive) cells at 24h and 48h after EdU addition during differentiation into neurons and osteoblasts as exemplarily shown for a region on chromosome 1 (Figure 5E) and a region on chromosome 12 (Figure 5F). In conclusion, the pattern of re-replicated DNA regions fits to the expression pattern.

**Figure 5:**
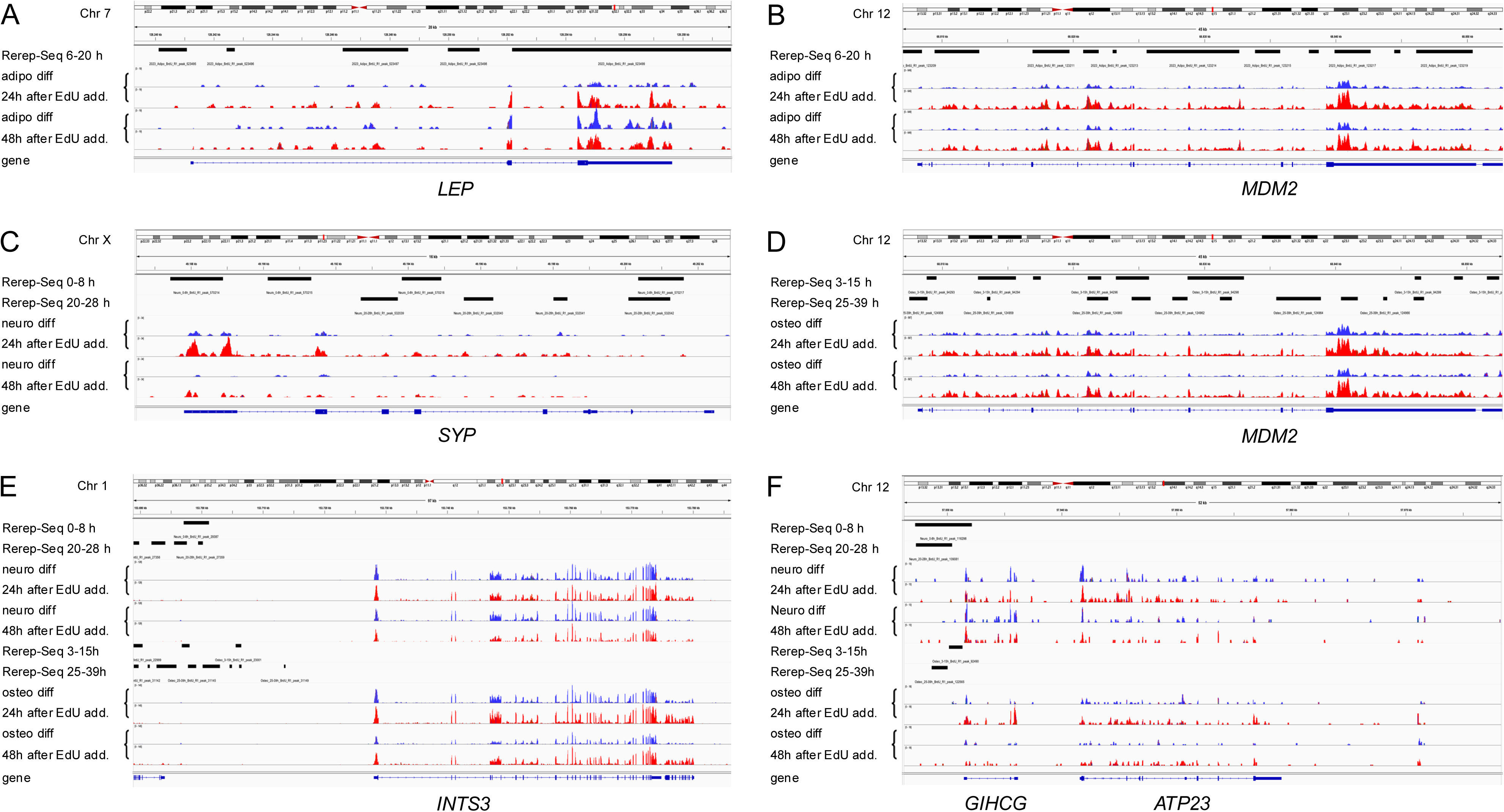
Expression analysis of re-replicated DNA Results of Rerep-Seq shown as broadPeaks were displayed with IGV together with results from RNA-Seq (24h and 48h after Edu addition for adipogenic differentiation (A, B), for neurogenic differentiation (C) for osteogenic differentiation (D) and for neurogenic and osteogenic differentiation (E, F). RNA-Seq results were represented by read-height displayed in red for replicating and in blue for not-replicating cells for *LEP* (A), *MDM2* (B and D), *SYP* (C), *INTS3* (E), and *GIHCG and ATP23* (F). BroadPeaks of re-replicated DNA overlap with increased read-height of genes *LEP*, *MDM2* and *SYP* at time points 24h and 48h post EdU addition in replicating (Edu-positive: red) versus not-replicating (EdU negative: blue) cells. Chromosome regions without broadPeaks of re-replicated DNA revealed similar read-height of genes *INTS3, ATP23* and lncRNA *GIHCG* in replicating (Edu-positive: red) and not-replicating (EdU negative: blue) cells 24h and 48h after addition of EdU.

### Exclusion of re-replicated DNA during proceeding differentiation

Previously, we reported evidence for clusters of amplified DNA outside the nucleus upon extended differentiation periods of myogenic differentiation (22). In the following, we analyzed whether DNA that is re-replicated during the differentiation processes examined here can also be detected outside of the cells. Following 3d and 4d of adipogenic differentiation and 3d of osteogenic differentiation, each in presence of EdU, hMSCs were fixed on glass slides and EdU was visualized with fluorescence dye Alexa-Fluor-594 (red). Extranuclear DNA spots and clusters were collected by microdissection. Only those red spots and clusters were collected that could also be identified by blue fluorescence (Hoechst33342-staining) of DNA to discriminate random fluorescence background spots from extranuclear DNA (Figure 6A). In total, 1,200-1,600 microdissected extranuclear DNAs were collected and sequenced. Notably, many enriched extrachromosomal DNA reads overlapped with re-replicated DNA stretches that were identified as broadPeaks in the differentiation assays described above including genes for which gene-amplification was previously reported. Examples of these enrichments were shown for *MDM2* (Figure 6B) and *LEP* (Figure 6C). In summary our data indicate that extranuclear DNA originates from DNA regions that have undergone re-replication.

**Figure 6:**
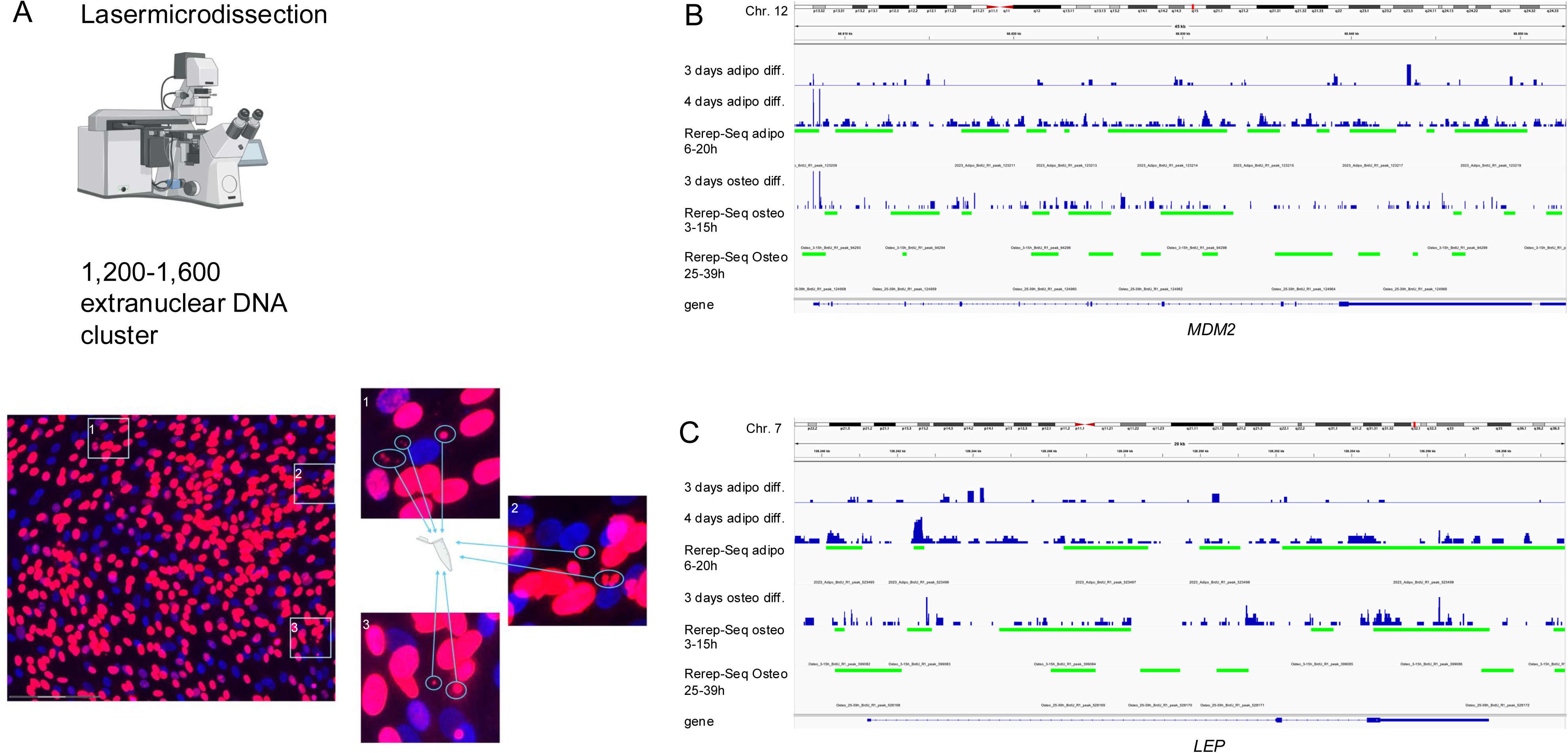
Analysis of extranuclear DNA during differentiation Mesenchymal stem cells were grown on glass slides with corresponding differentiation media and supplemented with thymidine-analogue EdU during timeframes of re-replication determined before. EdU incorporation is detected with red fluorescence (Alexa-Fluor-594) and DNA was counterstained with Hoechst33342 in blue. An example of osteogenic differentiated hMSCs is shown and three enlarged views point to extranuclear spots and clusters of DNAs that were microdissected (A). In total 1200-1600 extranuclear DNAs were collected for further DNA isolation and sequencing for each differentiation experiment. During adipogenic differentiation extranuclear DNA was detectable after 3d and 4d and during osteogenic differentiation after 4d of differentiation. Results of sequencing analysis of extranuclear DNA were displayed using IGV with data range on the y-axis adjusted for all extranuclear DNA reads. For each differentiation extranuclear DNA sequencing (dark blue) is displayed above the corresponding Rerep-seq results (green). Representative examples for gene regions on chromosome 12 (B) and 7 (C) revealed that extranuclear DNA overlaps with re-replicated DNA regions.

### Evidence for asymmetric re-replication

In addition to the exclusion of re-replicated DNA from the nucleus, re-replication may deviate from the proposed “onion-skin model,” as it poses a significant threat to genome stability. Analogous to findings in *Drosophila*, where Osheim et al. described asymmetric re-replication bubbles (23), our fiber-combing experiments revealed several replication forks, where re-replication was observed in various forms: occurring on two strands of the replication fork but differing in length between strands (Figure 7A), occurring only on one strand with no re-replication on the other strand of the replication fork (Figure 7B) and occurring only on one strand showing no detectable replication at the other strand of the replication fork (Figure 7C).

**Figure 7:**
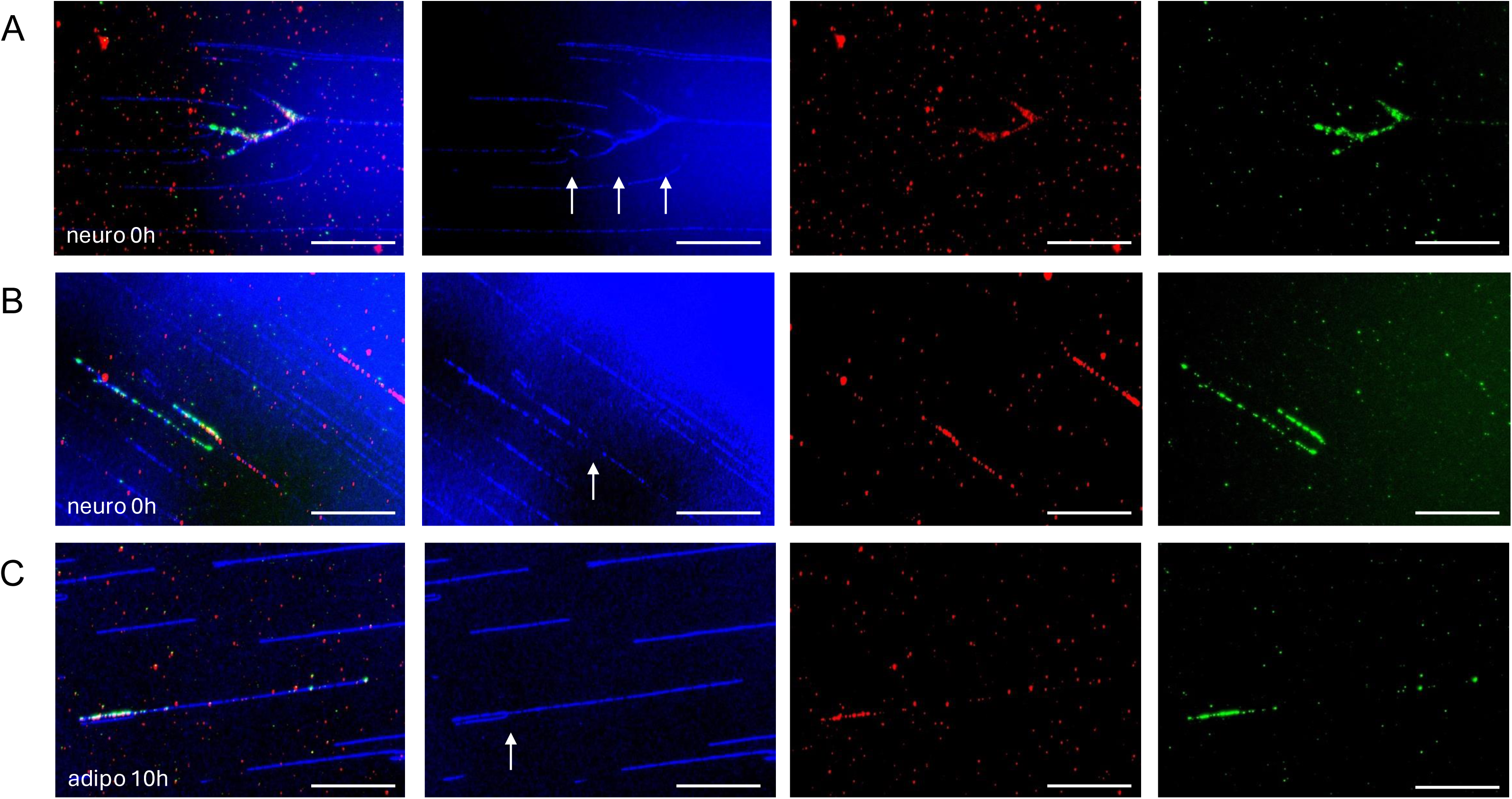
Asymmetric re-replication Fiber-combing experiments revealed several replication forks with varying combinations of replication and/or re-replication events. Representative examples shown, derived from neurogenic (A, B) and adipogenic (C) differentiated mesenchymal stem cells. Re-replication on two strands of the replication fork but differing in length between strands (A), re-replication only on one strand with no re-replication on the other strand of the replication fork (B) and re-replication only on one strand showing no detectable replication at the other strand of the replication fork (C). DNA is shown in blue (false color YOYO-stain) and thymidine-analogue detection is shown in red for IdU and green for CldU. Re-replicated fiber tracks appear as yellow fluorescence. Scale bars represent 20 µm = 40kb.

## DISCUSSION

Replication is typically restricted to once per cell cycle, but during development, this restriction is intentionally bypassed to enable endoreplication, a phenomenon well-studied in *Drosophila*. In this study, we demonstrate that re-replication also occurs during hMSC differentiation. In earlier studies, we demonstrated gene amplifications in hMSCs during differentiation into adipocytes or osteoblasts (6). The hypothesis that a re-replication mechanism underlies these amplifications is supported by comparative studies with other species. In *Saccharomyces cerevisiae*, it has been shown that re-replication can trigger gene amplification, with loss of replication control acting as a potent inducer of this process (3). Specifically, repression or deletion of the replication regulator Geminin (*GMNN*) induces re-replication (24), and overexpression of *CDT1*, which is normally degraded during the mitotic cycle by E3 ubiquitin ligation, also leads to re-replication (24). As addressed below, the homologs of these genes are also involved in replication control in human cells. The best-known example of physiological gene amplification is found in *Drosophila*. Endoreplication occurs in various cell types during *Drosophila* development, with endocycles and endoreplication proposed as key mechanisms driving this copy number gain. In mammalian cells induced overexpression of key replication regulators (*CDT1* and *CDC6*) or deletion of their repressors (*GMNN* or *Emi1*) causes re-replication (25–27).

Our evidence of physiological re-replication in hMSCs during differentiation into adipocytes, osteoblasts, chondrocytes, or neurons is based on two complementary approaches. Fiber-combing reveals extended stretches of DNA, indicating that a second replication event can begin before the first has been completed. While this method allows for visual single-molecule analysis, it has limitations: it does not specify where in the genome these re-replication events occur and captures only a small proportion of such events due to its focus on individual molecules. Additionally, regarding potential false positives in fiber-combing, there is a possibility that IdU-molecules remain, even after washing with PBS prior to adding CldU. In this scenario, both IdU and CldU could be incorporated into the DNA during replication, resulting in a yellow track without actual re-replication occurring. To minimize the risk of false positive re-replication events, we established a very stringent threshold of 10kb minimum length required to classify yellow tracks as re-replication event.

The second approach, termed Rerep-Seq, which was recently developed, is based on the enrichment of DNA fragments due to an increased number of strand breaks resulting from BrdU incorporation during re-replication events. While this approach allows the mapping of re-replication events to gene regions, it also has inherent drawbacks. First, Rerep-Seq may lead to false-positive results because it relies on the enrichment of small DNA fragments that are isolated and sequenced. Chromosomal regions in which breaks occur physiologically very frequently, such as fragile sites, may generate small DNA fragments without re-replication events. In addition, re-replication itself is accompanied by double strand breaks that generate fragments. Consistently, we identified weak DNA fragmentation in our control experiments without BrdU. However, such false-positive fragments can be excluded since they are generated both in the presence and absence of BrdU and were filtered out by the MACS2 peak calling tool. Secondly, Rerep-Seq can also lead to false-negative results since re-replication may be overlooked. Following BrdU incorporation, frequent re-replication events within a DNA region and an increase of double strand breaks can generate very small DNA fragments, which are too small to be detected by Rerep-Seq. This shortcoming must also be considered when interpreting the broadPeaks that indicate re-replication events. A gene region for which several, but no continuous, broadPeaks are observed may be continually re-replicated. Our experiments with an early and a later Rerep-seq timeframe indicated a stepwise complete coverage of the region. Nevertheless, Rerep-Seq allows for the identification and broad mapping of re-replication events to chromosomal regions.

The re-replicated regions identified here overlap with chromosomal regions previously shown to be amplified during differentiation, including *MDM2* gene (chromosome 12) and *PRSS1* gene (chromosome 7) during adipocyte and osteoblast differentiation (6). During myoblast differentiation the amplified *CDK4, MDM2* and *NUP133* genes overlap with re-replicated regions identified here and thereby complement our previous fiber-combing analysis on myotube differentiation (20, 22). This finding provides further evidence that re-replication acts as a mechanism for gene amplification in response to specific, time-limited demands for protein generation during stem cell differentiation.

This study and previous analyses show that not all cells within a differentiating stem cell population exhibit re-replication, gene amplifications, and concomitant overexpression of genes that map to re-replicated and amplified regions. This divergence may, in part, be due to bulk gene expression analyses. In the current study we enriched re-replicating cells by FACS sorting and identified overexpressed genes in these cells compared to non-replicating cells. Using the thymidine analog EdU, which can be easily detected with a fluorescent dye, we can separate replicating and non-replicating cells and study their specific mRNA expression. Since normal replication during cell division is negligible in differentiating cells (e.g. <0.5% for adipogenic differentiation), the replicating cells mostly correspond to re-replicating cells. These can be used to analyze the downstream effects of re-replication on corresponding mRNA expression. We chose 24 and 48h after EdU labeling to allow for sufficient transcription of re-replicated DNA. The overexpressed genes include those that play roles in specific differentiation processes, such as *SYP* (marker of synapse maturation) (14) overexpressed in re-replicating cells during neuronal differentiation. Another example is re-replication and overexpression of the adipocyte specific gene *LEP*. In addition to these examples of re-replicated and overexpressed genes we would like to point on another correlation. There were many chromosomal regions without any re-replication detected all over the genome and all differentiation experiments. When going into details of these regions many of them included lncRNAs with so far unknown regulatory functions including *GIHCG*, oncogenes or cancer associated genes including *AKT3* and *KIF26B* or double-strand-break-repair associated genes including *ATP23* and *INTS3*. These genes revealed similar expression in re-replicating versus not replicating cells. Although it looks like a random distribution of these not re-replicated regions, the content of the regions and above-mentioned genes and lncRNAs strongly argues for a selective repression of these genes. Since re-replication leads to double strand breaks, single strand breaks and an enhanced risk of genome instability it is reasonable to repress repair of this risky fragmented DNA and to repress oncogene functions.

However, re-replicating cells suffer a high risk of genome instability without removal/segregation of fragmented DNA. Beside reducing the risk of genome instability not re-replicating neighboring cells could benefit from this segregated DNA. Provided that the extranuclear DNA from re-replicating cells is taken up by not re-replicating cells, this would ensure those cells overexpression of proteins essential for differentiation. To address this hypothesis, we used laser-microdissection and isolated extranuclear DNA from adipogenic and osteogenic differentiated cells 3–4d post differentiation induction. Sequence analysis of segregated DNA revealed overlaps with re-replicated regions. Also consistent with this mechanism are early studies on leukemia, which demonstrated amplified *MYC* genes excluded from the nuclei and localized on double minutes (28). Since these amplified gene sequences, released from the cell nucleus, are also associated with an improved prognosis, they may also be functional.

Therefore, we propose a model in which only a portion of the cells undergo re-replication during stem cell differentiation. Cells that do not undergo re-replication avoid the risk of genome instability due to strand breaks associated with re-replication. In this model, re-replicated DNA is transferred from a re-replicating cell to a non-replicating cell. As a result, these non-replicating cells can ensure the specific increased production of proteins required during differentiation as summarized for differentiation of hMSCs to adipocytes in Figure 8.

**Figure 8:**
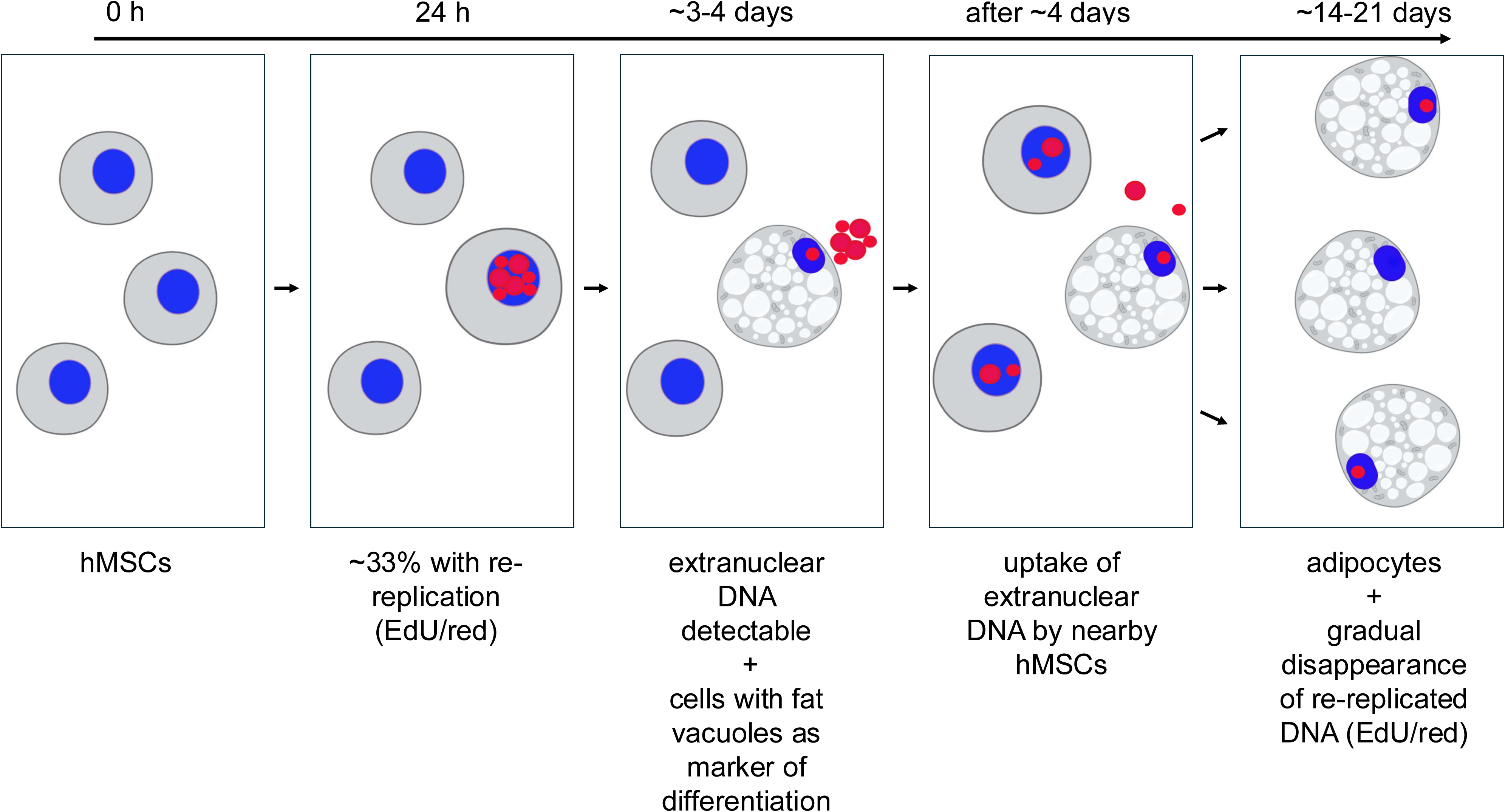
Postulated model of extranuclear DNA reuse during differentiation During differentiation exemplarily shown for adipocyte differentiation, only ∼1/3 of hMSC revealed replication (EdU incorporation, red fluorescence) after 24h, the remaining hMSCs revealed no replication and the nucleus is only stained blue with Hoechst33342. After 3-4d DNA with EdU was detectable outside the nucleus and outside the cells and cells revealed fat-vacuoles as sign of differentiation into adipocytes. We postulate that during proceeding differentiation extranuclear DNA is taken up by not re-replicating cells. Afterwards, more and more cells are differentiated into adipocytes and extranuclear DNA gradually disappears and thereby re-replicated DNA is no more a risk for genome instability. The figure was created using BioRender.

## ACKNOWLEDGEMENTS

We acknowledge the technical help of Esther Maldener.

## CONFLICT OF INTEREST

The authors have no relevant financial or non-financial interests to disclose.

## DATA AVAILABILITY

*The data underlying this article are available in* [SRA] [Bioproject accession number PRJNA1240097].

## AUTHOR CONTRIBUTIONS

M.M.: investigation, writing – review and editing; A.B.: formal analysis, writing – review and editing; S.R.: formal analysis, data curation, writing – review and editing; E.M.: investigation, writing – review and editing; D.Y.: investigation, visualization, writing –, review and editing; G.P.S.: formal analysis, data curation, writing – review and editing; P.E.S.: investigation, writing – review and editing; M.S.: investigation, writing – review and editing; T.T.: investigation; M.C.: investigation, writing – review and editing; N.L.: investigation, writing – review and editing; A.K.: conceptualization, funding acquisition, writing – review and editing, E.M.: conceptualization, funding acquisition, writing – original draft, review and editing, U.F.: conceptualization, investigation, visualization, writing – original draft, review and editing.

## FUNDING

This work was supported by Deutsche Forschungsgemeinschaft DFG INST 256/541-1 FUGG to E.M.; DFG INST 256/508-1 and DFG INST 256/550-1 FUGB to D.Y. The compute infrastructure for this project was funded by Deutsche Forschungsgemeinschaft DFG [469073465] to A.K. This work was further supported financially by Saarland University and the UdS-HIPS TANDEM initiative to A.K. and Funding “Anschubfinanzierung 2025” of Saarland University to U.F. Funding for open access charge: Saarland University.

## Notes

### Competing Interest Statement

The authors have declared no competing interest.

