## Supplementary Figure 1 and Supplementary Figure 2 for "Physiological re-replication during human stem cell differentiation"

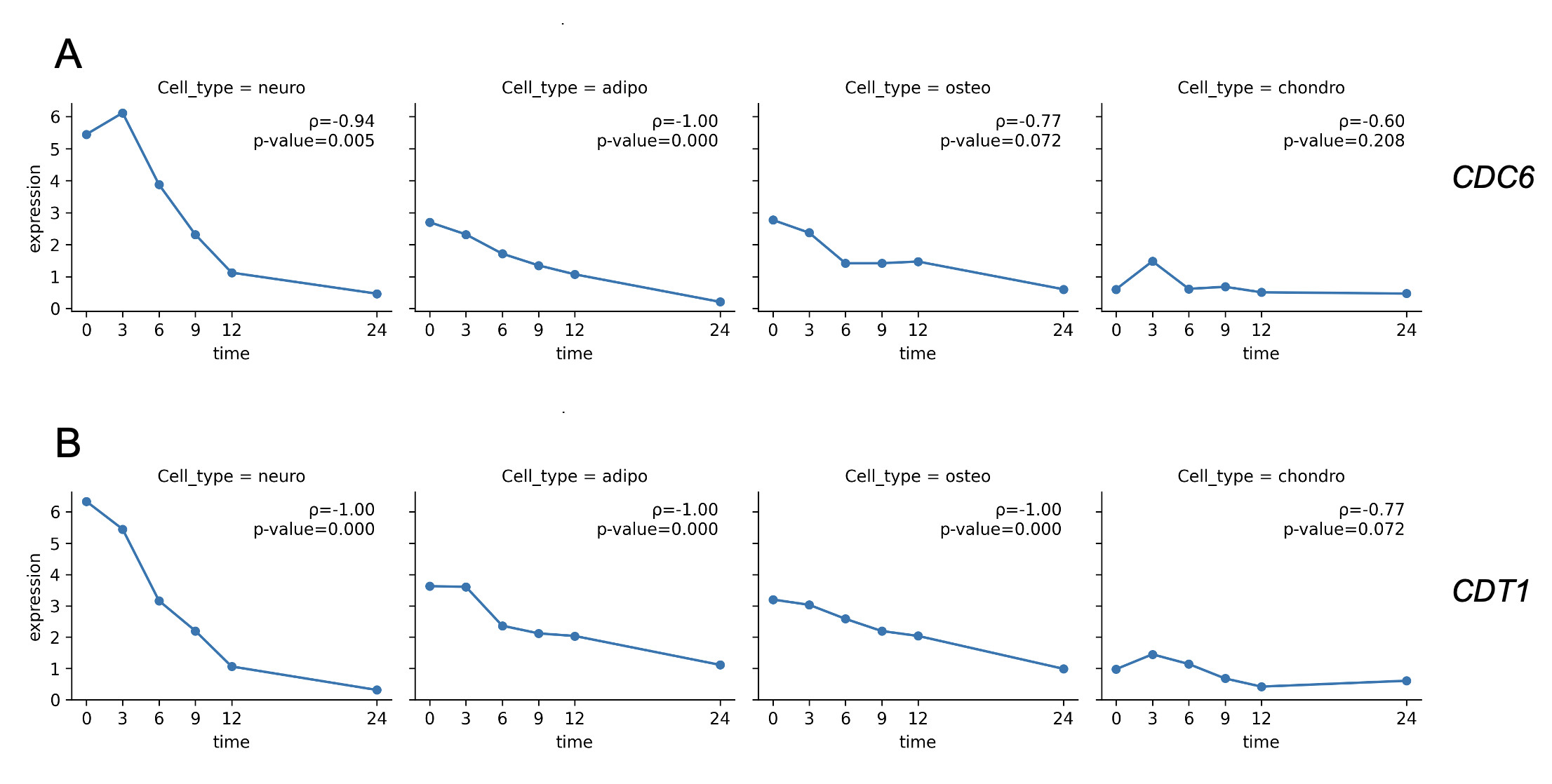

**Supplementary Figure S1: Expression analysis of *CDC6* and *CDT1***

Results of RNA-Seq of 24h neuron, adipocyte, osteoblast and chondrocyte differentiation were shown for *CDC6* in the upper part (A) and *CDT1* in the lower part (B). The graphs display the calculated spearman rank correlation coefficient and corresponding p-value for each gene expression (rpkm) with time (hours).

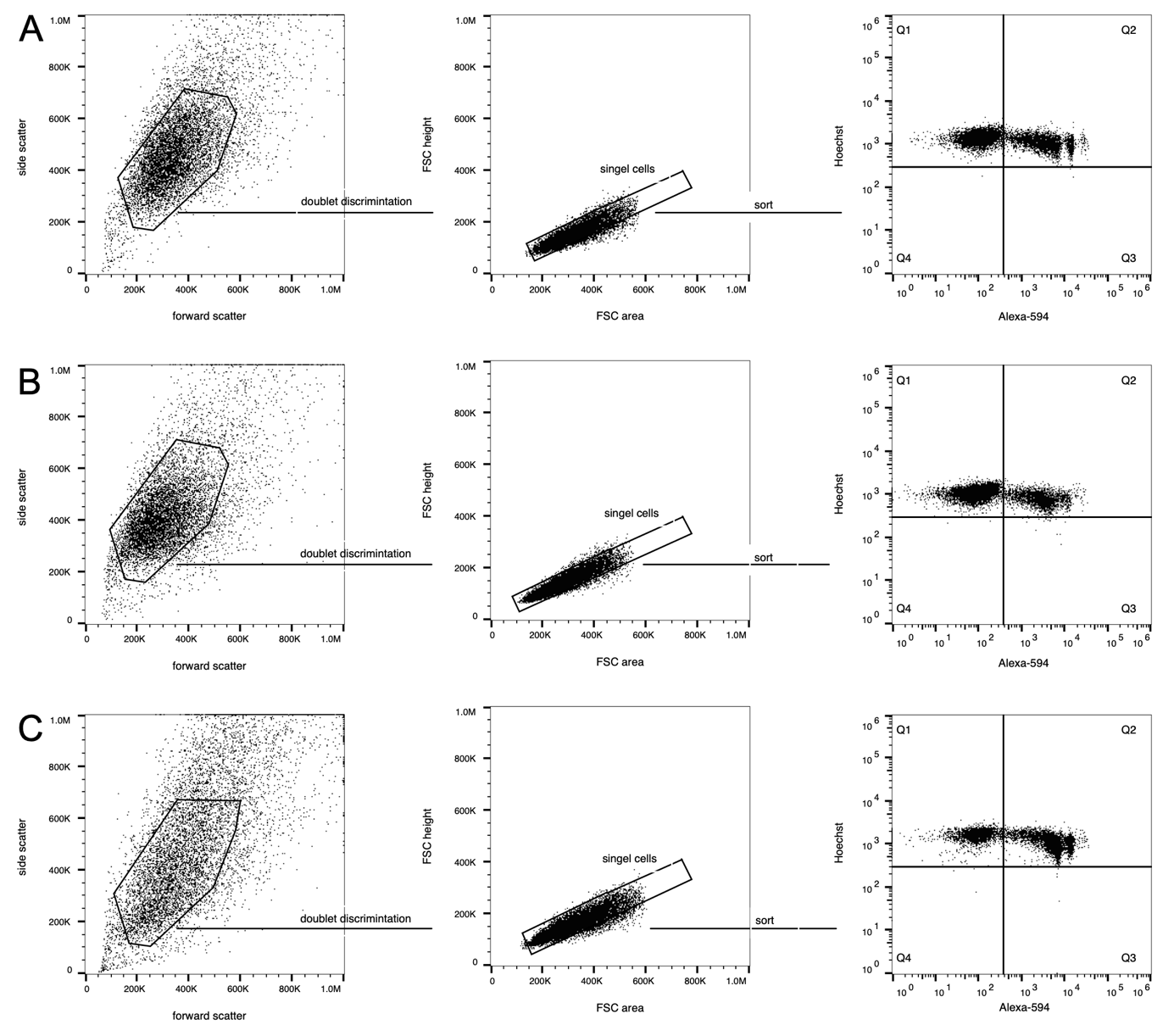

**Supplementary Figure S2: Gating strategy for cell sorting**

The homogenous cell population was identified and separated from cell debris by forward scatter (FSC) and side scatter (SSC), followed by FSC area versus FSC height discrimination of doublets. Unstained cells (without Hoechst 33342 and Alexa Fluor 594) und single stained cells were used to set the initial gates. Finally, cells were sorted by fluorescence with excitation at 405 nm (violet laser) and 638 nm (red laser), and emission was detected in FL1 (450/50 nm) and FL4 (665/30 nm). Quadrant 1 (Q1) contains cells without replication (Hoechst 33342^+^Alexa Fluor 594^-^), Q2 cells with replication (Hoechst 33342^+^Alexa Fluor 594^+^). The gating strategy is shown for adipocytic (A), neuronal (B), and osteogenic (C) differentiation.
